# Genetic Discovery Enabled by A Large Language Model

**DOI:** 10.1101/2023.11.09.566468

**Authors:** Tao Tu, Zhouqing Fang, Zhuanfen Cheng, Svetolik Spasic, Anil Palepu, Konstantina M. Stankovic, Vivek Natarajan, Gary Peltz

## Abstract

Artificial intelligence (AI) has been used in many areas of medicine, and recently large language models (LLMs) have shown potential utility for clinical applications. However, since we do not know if the use of LLMs can accelerate the pace of genetic discovery, we used data generated from mouse genetic models to investigate this possibility. We examined whether a recently developed specialized LLM (Med-PaLM 2) could analyze sets of candidate genes generated from analysis of murine models of biomedical traits. In response to free-text input, Med-PaLM 2 correctly identified the murine genes that contained experimentally verified causative genetic factors for six biomedical traits, which included susceptibility to diabetes and cataracts. Med-PaLM 2 was also able to analyze a list of genes with high impact alleles, which were identified by comparative analysis of murine genomic sequence data, and it identified a causative murine genetic factor for spontaneous hearing loss. Based upon this Med-PaLM 2 finding, a novel bigenic model for susceptibility to spontaneous hearing loss was developed. These results demonstrate Med-PaLM 2 can analyze gene-phenotype relationships and generate novel hypotheses, which can facilitate genetic discovery.

## 1. Introduction

A major challenge in biomedical science is to identify genetic factors in a population that affect the properties (i.e., phenotypes or traits) of an individual, especially those for disease susceptibility. Many genetic discoveries have been made using genome wide association study (GWAS) methods, which compare the pattern of allelic differences in mouse or human populations with variation in phenotypic responses. Irrespective of whether the subjects are mouse or human, a major barrier for genetic discovery is that GWAS results will identify a true causative genetic variant along with multiple other false positive associations because allelic patterns within genomic regions that are commonly inherited from their ancestors can randomly correlate with any phenotypic response pattern in a population. To address this issue, we recently developed an artificial intelligence (AI)-based computational pipeline for mouse genetic discovery that could sift through a set of candidate genes emerging from a GWAS and identify those most likely to be causal based upon assessment of the candidate gene-phenotype relationships in the published literature and other factors [1]. This pipeline analyzed publicly available datasets of biomedical responses (or disease susceptibility) that were measured in panels of inbred strains and identified novel causative genetic factors for various disease models, which included diabetes/obesity and cataract formation. However, this pipeline could only analyze a specific dataset and the analysis required entry of its numeric label.

The utility of a genetic discovery AI would be greatly expanded if it could answer free-text queries. Consistent with this possibility, large language models (LLMs) have recently been developed, which are general-purpose AI systems that are commonly implemented with transformer based neural network architectures [2]. They are typically pretrained on internet scale text corpora [3] and have demonstrated a wide range of language understanding and processing capabilities including information extraction, reasoning, summarizing data, and entailment capabilities. While general-purpose LLMs encode knowledge and have demonstrated their capabilities across a wide range of applications, the uniqueness of the scientific and biomedical domain requires that they be further specialized and adapted. LLM task performance can be improved through fine-tuning, which uses high quality domain-specific texts (i.e., scientific and biomedical text, or clinical guidelines) and/or learning from expert feedback. For example, Med-PaLM 2 is a recently developed medically aligned LLM that was fine-tuned using high quality biomedical text corpora and was aligned using clinician feedback. It demonstrated significantly improved performance when answering medical questions relative to general-purpose LLMs, and its answers matched or exceeded those of physicians when controlled benchmarks were assessed [4, 5]. Despite these advances and the large volume of biomedical and scientific knowledge encoded within LLMs, it remains to be determined if LLMs can be used to generate novel hypotheses that facilitate genetic discovery. Here, we test the potential of the specialized Med-PaLM 2 model to uncover gene-phenotype associations (Figure 1). We found that Med-PaLM 2 correctly responded to free-text queries about potential sets of candidate genes and that it could identify a novel causative genetic factor for an important biomedical trait.

**Figure 1.**
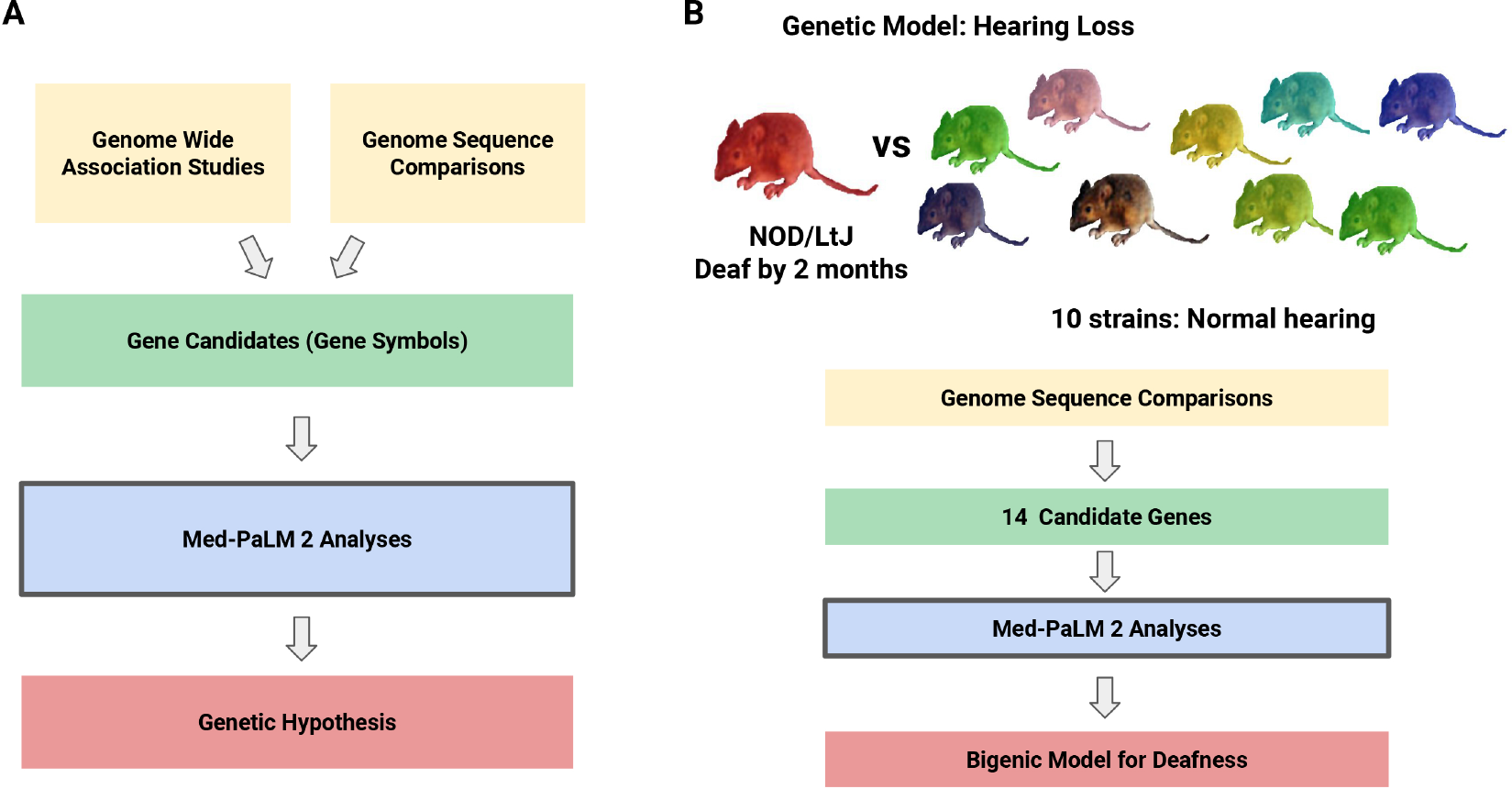
The Med-PaLM 2 pipeline for genetic discovery. **(A)** Overview of the genetic analysis pipeline. A set of candidate genes are identified through analysis of the results obtained from either Genome Wide Association Study (GWAS) or genomic sequence comparison. Med-PaLM 2 evaluates the gene candidates (represented by their gene symbols) and it generates a genetic hypothesis by identifying those with the strongest association with a queried phenotype. **(B)** Overview of how Med-PaLM 2 generated a bigenic model for spontaneous hearing loss in a mouse strain. The genomic sequence of a mouse strain (NOD/LtJ) that spontaneously develops a hearing loss by 7 weeks of age was compared with that of 10 strains that maintain normal hearing during their lifetime. Fourteen genes with NOD/LtJ-specific high impact alleles were identified by this analysis. Med-PaLM 2 identified one gene as the most likely to contain the causative genetic factor for hearing loss. However, NOD/LtJ mice have another genetic factor, that is shared among multiple inbred strains with early onset hearing loss, which is necessary but it alone is not sufficient to cause their severe hearing loss. Therefore, based upon the Med-PaLM 2 results, the genetic hypothesis developed was that two genetic factors (i.e., a bigenic model) jointly contribute to the hearing loss of NOD/LtJ mice.

## 2. Results

### Assessing candidate gene sets

To investigate whether Med-PaLM 2 could be used for genetic discovery in response to free-text input, it was asked to analyze sets of candidate genes, which were previously identified by analysis of mouse GWAS data for six previously studied biomedical traits and identify the most likely causative genetic factors. In each example, Med-PaLM 2 was asked to identify which gene among a set of input genes was most likely to be associated with a queried phenotype. The candidate gene sets for cataract and diabetes susceptibility were identified using the mouse genetic discovery AI [1]; and those for the albino skin color [6], warfarin metabolism [7], aromatic hydrocarbon response [6], and chronic pain [8] phenotypes output by the computational genetic mapping program after analysis of the data obtained from inbred mouse strain panels. For all six examples, Med-PaLM 2 correctly identified the gene with the experimentally verified causative factor (Table 1, Table ED.1 and Table ED.2).

**Table 1.**
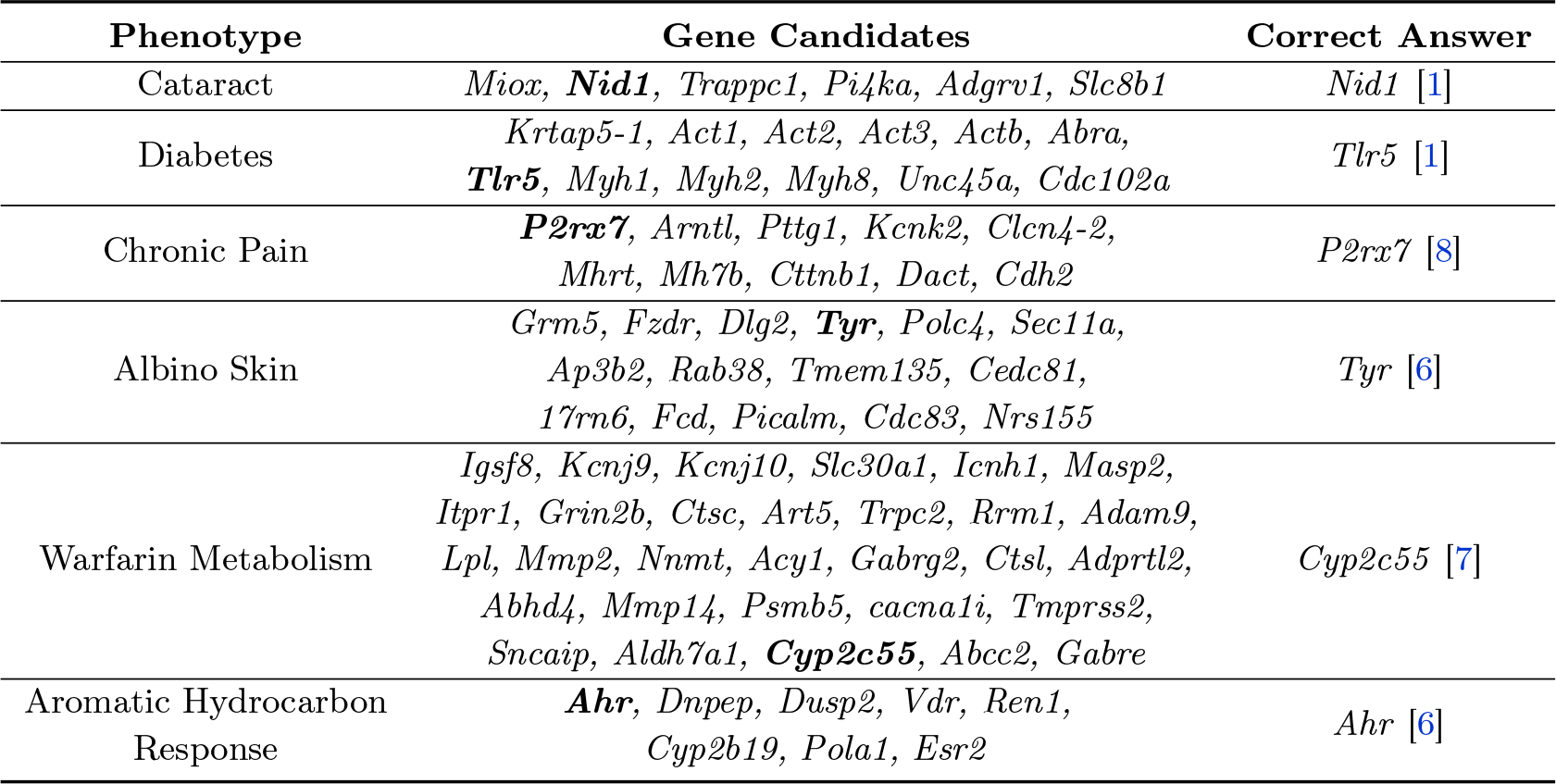
This table shows the candidate genes sets analyzed by Med-PaLM 2. The murine candidate genes, which had allelic patterns that correlated with the pattern of phenotypic responses of the inbred strains for the indicated phenotype, for each referenced study are listed. The experimentally proven causative genetic factor, which was correctly identified by the Med-PaLM 2 response is in bold. The queries and responses are shown in Table ED.1 and Table ED.2.

While Med-PaLM 2 correctly identified the experimentally verified gene (*P2rx7*) [8] in response to the chronic pain query, it also mentioned two other genes whose potential relationship to chronic pain we had not previously considered: (i) *Kcnk2* is a potassium channel that affects multiple types of pain responses [9–11]; and (ii) *Arntl*, which is a major circadian clock regulator [12], because maladaptive pain is affected by circadian rhythm disruption [13]. Although the connection between *Arntl* and chronic pain is tenuous, the hypothesis that *Kcnk2* alleles could impact chronic pain is a creative and intriguing one. These results indicate that Med-PaLM 2 can assess a list of genes and identify those most likely to contain genetic factors for a studied trait, and also that it could generate novel hypothesis about gene-phenotype relationships.

### A bigenic model for hearing loss

We next investigated whether Med-PaLM 2 could facilitate novel genetic discovery by determining if it could identify murine genes with genetic variants, which were identified by genomic sequence comparison, that were most likely to cause hearing loss. Sixteen inbred mouse strains (of 80 tested) spontaneously developed an age-related nonsyndromic hearing loss by 3 months of age [14]. A *Cadherin 23* (*Cdh23 G->A*) frameshift allele [15] that reduced the stability of a cochlear sensory hair cell bundle protein was previously shown to contribute to hearing loss in mice [16, 17]. We examined our 25M SNP database with alleles covering 48 classical inbred strains [18, 19] and found an extremely strong correlation between strains with early onset hearing loss and *Cdh23*^*753A*^ alleles (Table SI.1). However, since only a subset of the strains with *Cdh23*^*753A*^ alleles developed early onset hearing loss, there must be other contributing genetic factors. The age-related hearing loss of NOD/LtJ mice was of interest because it occurs by 3 weeks of age [14] and we found that it is more severe than was previously known. NOD/LtJ cochlear sensitivity was dramatically reduced across all frequencies tested. C57BL/6 mice, which also have *Cdh23*^*753A*^ alleles, develop a hearing loss that is much less severe at 7 weeks of age (see Figure 2) and it is more fully manifested at a later age [14]. These features confirm that genetic factors other than *Cdh23*^*753A*^ alleles must contribute to the NOD/LtJ hearing loss, but none have yet been identified. To identify them, the genomic sequence of NOD/LtJ was compared with 10 other strains (A/HeJ, AKR, BALB/C, CBA, FVB, LG/J, PL/J, SJL, SM/J, SWR) that maintained normal hearing throughout much of their life [14], and variant alleles that were also present in any these other strains were removed. Among the high impact NOD/LtJ-specific SNP alleles (Table ED.4), those within genes that encoded polymorphic genomic markers, immunoglobulin variable region genes, or affected splice sites were removed. Med-PaLM 2 was then asked to analyze the remaining 14 genes using CoT prompting with self-consistency, and it identified *Crystallin mu* (*Crym*) as the gene that was most likely to be associated with hearing loss (Table ED.3). When Med-PaLM 2 analyses were repeated 100 times after the input gene order was randomized, *Crym* was identified as one of the top 5 candidates in 98 of the analyses (Table SI.2). NOD/LtJ mice have a 2-bp frameshift deletion allele (rs216145143) within the codon for amino acid 220 in *Crym* that was not present in 47 classic inbred strains [18, 20] including the closely related NOR/LtJ stain [21], which generates a truncated protein with a premature termination codon at position 230 (Figure 3A-C). Immunoblotting indicated that the CRYM protein is completely absent from NOD/LtJ tissue (Figure 3D), which results from degradation of mutated *Crym* protein or of its mRNA. Hence, the NOD/LtJ frameshift deletion makes NOD/LtJ mice the equivalent of a *Crym* knockout mouse.

**Figure 2.**
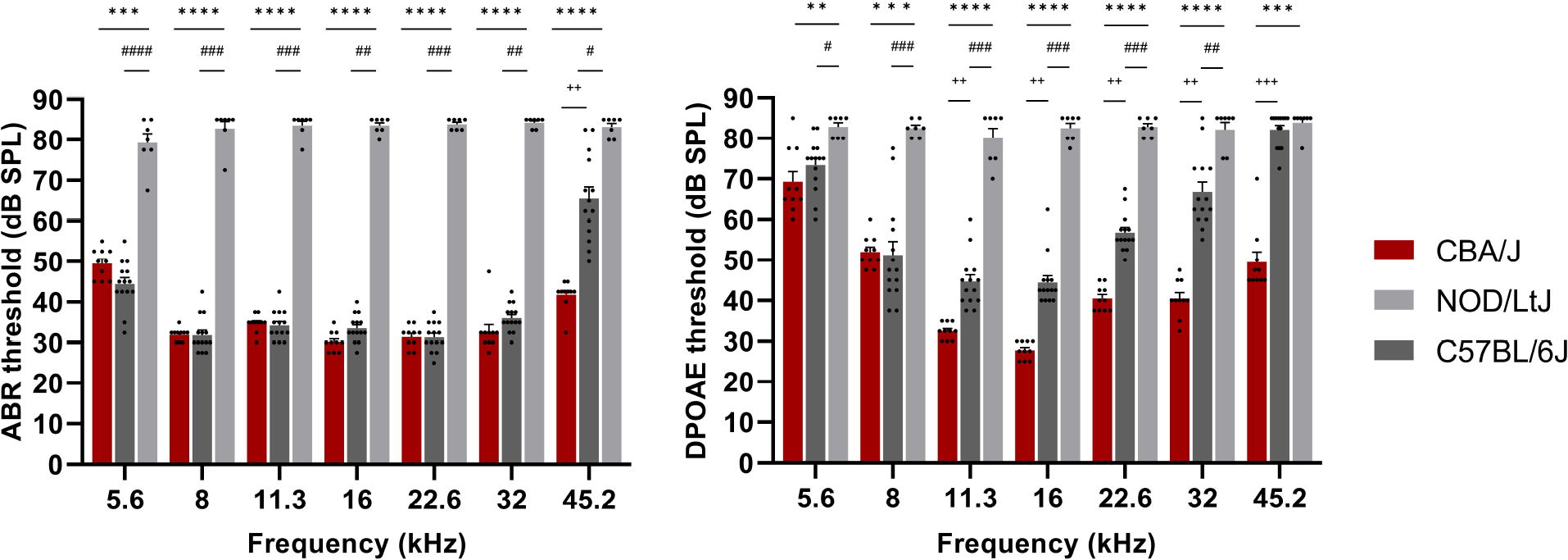
NOD/LtJ mice have a severe hearing loss. Auditory brainstem responses (ABR) and distortion product otoacoustic emissions (DPOAE) were measured in 7-week-old CBA/J (n=10, red), NOD/LtJ (n=7, light gray) and C57BL/6J (n=14, dark gray) mice. Each bar represents the mean ± standard error of the mean, and each dot represents the sound pressure level (SPL) in decibels measured for one mouse. The ABR threshold levels for NOD/LtJ mice demonstrate that they have a profound hearing loss compared to that of CBA/J and C57BL/6J mice of the same age. The DPOAE thresholds in NOD/LtJ mice are substantially elevated across all frequencies tested in comparison to that of CBA/J and C57BL/6 mice. Interestingly, C57BL/6 have a hearing defect that is less severe than that of NOD/LtJ mice, and they show significantly higher DPOAE thresholds in mid-to-higher frequency range than CBA/J mice. The p-values for the CBA/J vs. NOD/LtJ comparisons are represented by asterisks: *p<0.05, **p<0.01, ***p<0.001, and ****p<0.0001. Similarly, the p-values for C57BL/6J vs. NOD/LtJ comparisons represented by #, and + are used to represent the p-values for CBA/J vs. C57BL/6J comparisons.

**Figure 3.**
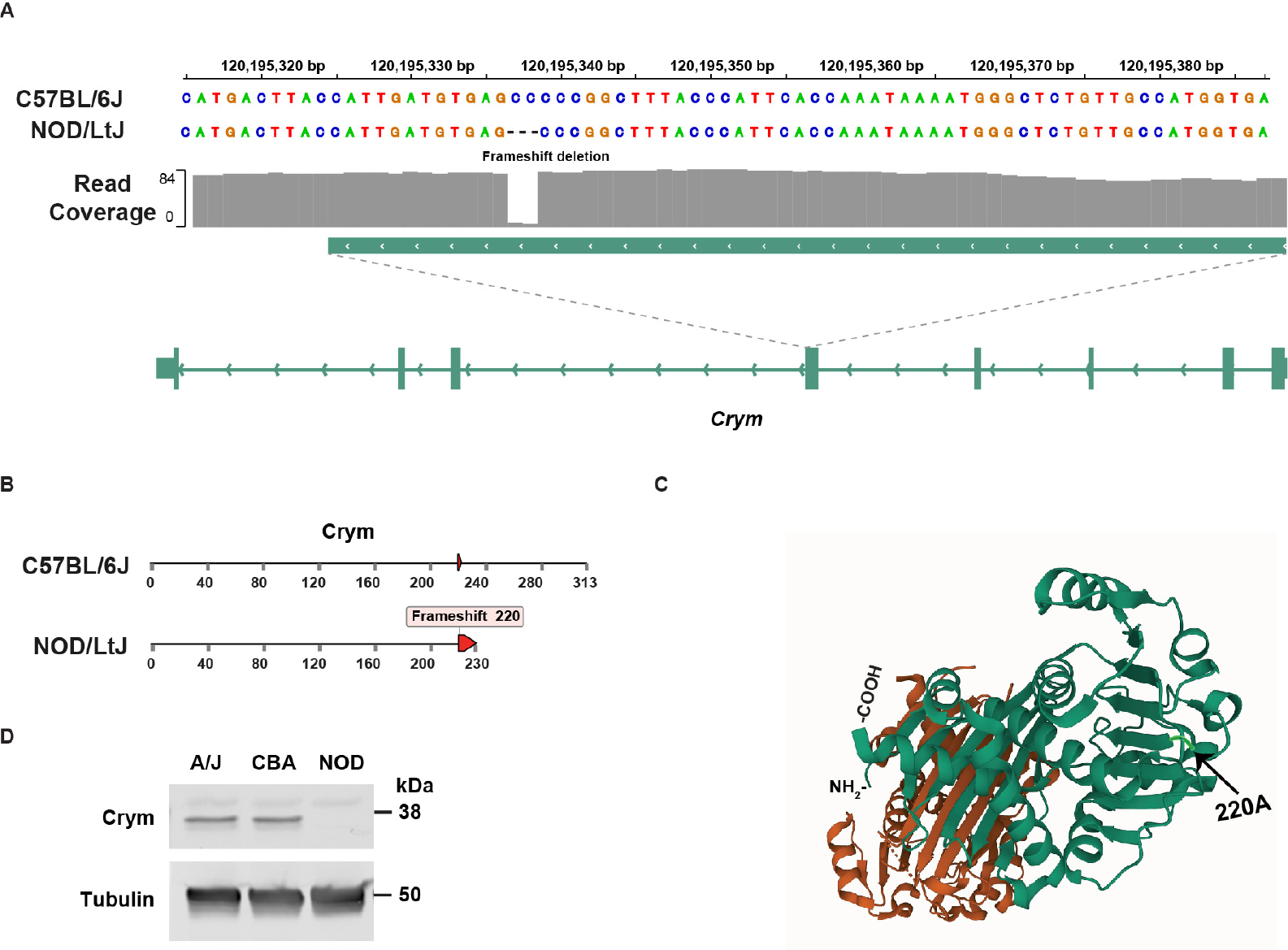
**(A)** NOD/LtJ mice have a 2 bp frameshift deletion in exon 4 of *Crym*, which is not present in 42 other strains. **(B)** The full length Crym protein has 313 amino acids, but the NOD/LTJ frameshift deletion within the codon for amino acid 220, which generates a premature termination codon at amino acid 230. **(C)** The Crym protein structure (PDB: 4BVA) is shown; and the position of NOD/LtJ-unique frameshift mutation that disrupts the COOH-terminal region of Crym is highlighted along with the location of the NH2- and COOH-terminal amino acids. **(D)** Crym protein is not present in NOD/LtJ mice. The proteins in lysates prepared from brain tissue obtained from a 5-week-old male A/J, CBA/J or NOD/LtJ mice were separated by SDS-PAGE and immunoblotted with mouse monoclonal anti-Crym or anti-tubulin antibodies. The blots were scanned after incubation with dye-labelled goat anti-mouse IgG. While the lysates have comparable amounts of tubulin, Crym is not present in the NOD/LtJ lysates.

*Crym* was identified by Med-PaLM 2 because it is highly expressed in a gradient (apex higher than base) along the length of the mouse cochlea [22], and two point mutations in human *CRYM* (*X315Y* or *K314T*) cause autosomal dominant non-syndromic deafness (DFNA40) [23, 24]. Thyroid hormone activity is essential for auditory function [25, 26]; CRYM is a triiodothyronine (T3) binding protein that regulates the intracellular level of T3 in cochlear cells [24] and the *K314T* mutation in human *CRYM* was shown to prevent thyroid hormone binding to CRYM [24]. Moreover, analysis of the crystal structure of Crym reveals that T3 binding is highly dependent upon the COOH-terminal domain of Crym: T3 binds to the aliphatic side chain of Arg229, Ser228 is one of two amino acids that hydrogen bonds with T3, and Ser228 and Leu292 interact with T3 through water molecules [27]. Since point mutations within the COOH terminal region of human CRYM cause early onset hearing loss, a NOD/LtJ *Crym* mutation that abolishes Crym protein expression is highly likely to contribute to its hearing loss.

## 3. Discussion

Based upon Med-PaLM 2 output, we propose that the spontaneous hearing loss of NOD/LtJ mice is likely to have (at minimum) a bigenic origin: the NOD/LtJ *Cdh23* and *Crym* frameshift alleles jointly contribute to it. This bigenic model explains four features of the hearing loss that develops in NOD/LtJ and in several other inbred strains. (i) Some inbred strains with *Cdh23*^*753A*^ alleles do not develop a hearing loss. (ii) The time of onset and the severity of the hearing loss developing in strains with *Cdh23*^*753A*^ alleles is variable. For example, the NOD/LtJ hearing loss occurs earlier and is more severe than in C57BL/6 mice, which have the *Cdh23*^*753A*^ allele but do not have the NOD/LtJ *Crym* frameshift allele. (iii) Reversion of the *Cdh23*^*753A*^ allele to wildtype only partially suppressed the hearing loss that develops in NOD mice [28]. (iv) *Crym* knockout mice were reported to have normal hearing [29], but this is a result of the *Crym* knockout being bred onto the 129Sv background: 129Sv substrains maintain normal hearing [14] throughout their life, and they have the *Cdh23*^*753G*^ that is associated with normal hearing. Since hearing loss does not develop in all strains with *Cdh23*^*753A*^ susceptibility alleles, the variability in the severity of the hearing loss in the strains with the *Cdh23*^*753A*^ susceptibility alleles, the inability of the reversion of the *Cdh23*^*753A*^ allele to wildtype in NOD mice to completely reverse the hearing defect, and the absence of a hearing loss in *Crym* knockout mice indicate that at least two genetic factors contribute to the NOD/LtJ hearing loss.

The Med-PaLM 2 analyses for hearing loss in a mouse is analogous to the methods used to identify causal genetic factors in newborns presenting with clinical abnormalities. Every year, around 7M newborns are affected by severe genetic diseases [30] and many are monogenetic diseases (MGD). Despite the increased use of genome sequencing, identification of MGD-causing variants is difficult because hundreds of rare variants of unknown significance (VUS) are present within many genes in any individual’s genome [31–35]. As in the mouse, the large number of VUS make it difficult to identify the true causative variant for a phenotype in a human subject’s genome. The magnitude of this problem is compounded by the fact that ∼ 60M suspected MGD patients will be sequenced by 2025. Numerous computational tools that automate some aspects of the MGD analysis pipeline have been developed to assess the potential pathogenicity of VUS [36–42]. Nevertheless, the rate of identifying disease-causing genes for suspected MGD (∼ 30%) is only modestly better than that achieved a decade ago [35]. Since Med-PaLM 2 can help identify disease-causing genetic variants in mice, it is likely that it can similarly be used to analyze VUS in human subjects.

Our findings also indicate how LLM genetic discovery performance can be improved. First, by analyzing the output, we identified at least two factors that contribute to the false associations generated by an LLM (which are referred to as ‘hallucinations’). (i) The tokenizer (i.e., the program that breaks down a piece of text into tokens) has difficulty analyzing gene symbols, which consist of letters (that are often acronyms) and numbers. Tokenizers can incorrectly assign gene symbols to other genes (or words), which generate false associations based upon analysis of the related genes (or words). For example, one input gene (*Med1*) that was not associated with hearing loss was associated with hearing loss because mutations in another gene (*MED12*) cause congenital hearing impairment [43]. (ii) The next word prediction objective used to train the LLM sometimes leads to spurious associations between genes and phenotypes, which do not have causal relationships, due to their co-occurrence within the same context window. In one case, a spurious association was caused by a series of >40 abstracts, each covering a different topic, that were published together as a conference summary. In another case, a diverse set of phenotypes exhibited by an inbred strain (which included hearing loss) were summarized in the introduction to a paper, which characterized the effect of a gene knockout on an unrelated phenotype [44]. Secondly, a scalable LLM based AI system for genetic discovery can be designed to overcome the current limitations of the knowledge encoded in LLMs. For example, such an AI system could be taught to use external tools, which include: advanced variant calling tools like AlphaMissense [45] to identify the more probable disease-causing alleles; or search based retrieval augmentation to provide supporting evidence (from other genetic data or the published literature) and remove false positive associations. Also, by using high-quality domain specific data, process supervision and providing granular feedback to the AI system, we can further enable the AI system to analyze regulatory elements, protein-protein interactions, or non-coding RNA functions which in turn will further expand the range of genetic hypothesis that these models can develop, and therefore lead to a deeper understanding of genetic diseases.

## Methods

### Animals

and Experimental Design. All mouse procedures were approved by the Institutional Animal Care and Use Committee at the Stanford University Medical School; they were conducted in accordance with the National Institute of Health Guide for Care and Use of Laboratory Animals, Eighth Edition; and the results are reported according to the ARRIVE guidelines [46]. Male NOD/LtJ, C57BL/6 and CBA/J mice were purchased from Jackson Laboratories (Sacramento, CA, United States). CBA/J mice were used as a control since they maintain good cochlear sensitivity through most of their adulthood. Cochlear function was examined by measuring auditory brainstem responses (ABR) and distortion product otoacoustic emissions (DPOAE) at 7 weeks of age (N=7 for NOD/LtJ, N=14 for C57BL/6, and N=10 for CBA/J).

### Cochlear function testing

Cochlear function testing was performed using previously described methods [47, 48]. In brief, mice were anesthetized with ketamine (120 mg/kg) and xylazine (12 mg/kg) that was administered intraperitoneally. A custom acoustic system consisting of two speakers (MF1, TDT, Alachua, FL, United States) and a microphone (ER-10B+, Ethymotic Research, Elk Grove Village, IL, United States) coupled to a probe tube was used to measure pressure near the mouse eardrum. DPOAEs were measured as ear canal pressure in response to two tones presented into the ear canal (f1 and f2, with f2/f1=1.2) at half-octave steps, from f2 = 5.6 to 45.2 kHz, and in 5 dB intensity increments from 10 to 80 dB sound pressure level (SPL). ABR responses to 5-ms tone pips were measured between subdermal electrodes (placed adjacent to ipsilateral pinna, at the vertex, and near the tail), amplified 10,000 times through an amplifier (Medusa4Z, TDT, Alachua, FL, United States) and filtered (0.3-3.0 kHz). For each frequency and sound level assessed, 512 responses were recorded and averaged using the BioSigRZ software run on a RZ6 Multi-I/O Processor (TDT, Alachua, FL, United States). ABR waveforms stacked from lowest to highest SPL were visually inspected to define the threshold as the first level at which a repeatable wave I was detected. In the absence of an ABR or DPOAE response, 85 dB SPL was chosen as a threshold because it was 5 dB above the highest sound pressure level tested [49].

### Whole genome variant calling and annotation

The pipeline used to identify and annotate the SNPs and structural variants in the database was previously described [1]. In summary, the raw reads of the genomic sequence for 53 inbred strains was aligned to the reference genome (GRCm38) using bwa [50] and variants were discovered using BCFtools [51]. Each variant was annotated using Ensemble Variant Effect Predictor (VEP) [52]. For this analysis, a variant was retained if it had a high-impact consequence on any transcript, as determined by VEP. To identify unique high-impact variants in the NOD/LtJ strain, we subtracted variant alleles that were also present in any of 10 other strains (A/HeJ, AKR, BALB/C, CBA, FVB, LG/J, PL/J, SJL, SM/J, SWR) that maintained normal hearing throughout much of their life [14]. VEP defines high-impact variants as those causing transcript ablation, splice acceptor/donor variants, stop gain or stop loss, frameshifts, start lost, or transcript amplification (Table ED.4). Alleles within genes encoding polymorphic genomic markers or immunoglobulin variable regions, or within splice sites were removed. Then, the 14 genes with alleles that caused a frameshift, stop loss, or stop gain were selected for analysis by Med-PaLM 2.

### Prompting strategies for Med-PaLM 2

To assess the capability of Med-PaLM 2 to identify a causative genetic factor for previously studied biomedical traits, Med-PaLM 2 was zero-shot prompted to select the most likely causative gene from a set of genes that were identified previously through computational genetic analysis of mouse GWAS data. We evaluated the model’s answers for the six biomedical traits because the correct causative gene for each phenotype was validated in previous studies. The prompts and model outputs for all 6 test examples are shown in Table ED.1 and Table ED.2.

To assess the ability of Med-PaLM 2 to facilitate novel genetic discovery, Med-PaLM 2 was used to identify murine genes with genetic variants that contribute to hearing loss. A set of 14 high impact NOD/LtJ-specific SNP alleles were identified as described above. Chain-of-Thought (CoT) and self-consistency prompting strategies were used to ask Med-PaLM 2 to output the top 5 most likely genes that could be associated with hearing loss. CoT involves augmenting a one-shot example in a prompt with a step-by-step explanation of how to move towards a final answer [53]. CoT prompts were crafted to provide clear demonstrations on how to appropriately answer the gene-phenotype association questions shown in Table ED.3. Since the problem was formulated as a multiple-choice question, a self-consistency strategy was used to determine the most consistent answer [54]. Specifically, 100 decoding outputs from the model were sampled by randomizing the input order of the candidate genes. At each inference time, Med-PaLM 2 was instructed to select the top 5 genes associated with hearing loss. The total count for the number of times that each gene was selected across 100 decodes was aggregated.

### Immunoblotting

Brain tissues obtained from a 5-week-old male A/J, CBA/J or NOD/LtJ mice were homogenized in RIPA buffer supplemented with a protease inhibitor cocktail (Sigma P8340, 1 to 100) using a Precellys tissue homogenizer. Thirty micrograms each of protein lysate were resolved on a 4-15% gradient SDS-polyacrylamide gel and transferred to a nitrocellulose membrane. The membranes were incubated with a 1:1000 dilution of mouse monoclonal anti-*µ*-crystallin antibody (F-11) (Santa Cruz Biotechnology) or with a 1:4000 dilution of a mouse monoclonal anti-*α*-tubulin antibody (clone B 5-1-2) (Sigma Aldrich). The anti-Crym antibody is specific for an epitope that is located between amino acids 45 and 85 of the Crym protein. The membranes were then incubated with IRDye^@^ 800CW goat anti-mouse IgG (Licor Bioscience) as the labeled secondary antibody, and were scanned using a Licor Odyssey imaging system.

## Acknowledgments

This work was supported by NIH awards (1R01DC021133 and 1 R24 OD035408) to GP. TT, VN, and AP were funded by Alphabet Inc. and/or a subsidiary thereof (Alphabet). We thank Pi-Chuan Chang and Greg Corrado for their valuable insights and feedback during our research.

## Data Availability

SNP allele data is available at the Mouse Phenome Database (GenomeMUSter https://mpd.jax.org/genotypes).

## Code Availability

Med-PaLM 2 is an LLM for the biomedical domain. We are not open-sourcing model code and weights owing to the safety implications of unmonitored use of such a model in medical settings. In the interest of responsible innovation, we will be working with academic and industry research partners, providers, regulators, and policy stakeholders to validate and explore safe onward uses of Med-PaLM 2. For reproducibility, we documented technical deep learning methods while keeping the paper accessible to a clinical and general scientific audience. Our work builds upon PaLM, for which technical details have been described extensively, and our institution has open-sourced several related LLMs to further the development of research methods in the field (https://huggingface.co/google/flan-t5-xl).

## Author contributions

GP and KS formulated the project. TT and GP wrote the paper with input from all authors. TT, FZ, SS, and ZC generated experimental data. TT, FZ, and GP analyzed the data. TT, AP, and VN developed the LLM and the techniques for enabling its genetic discovery applications. All authors have read and approved of the manuscript.

## Competing interests

The Stanford University Medical School authors have no competing interests. T.T., V.N., and A.P. are employees of Alphabet and may own stock as part of their standard compensation package.

## Supplementary Information

**Table SI.1.**
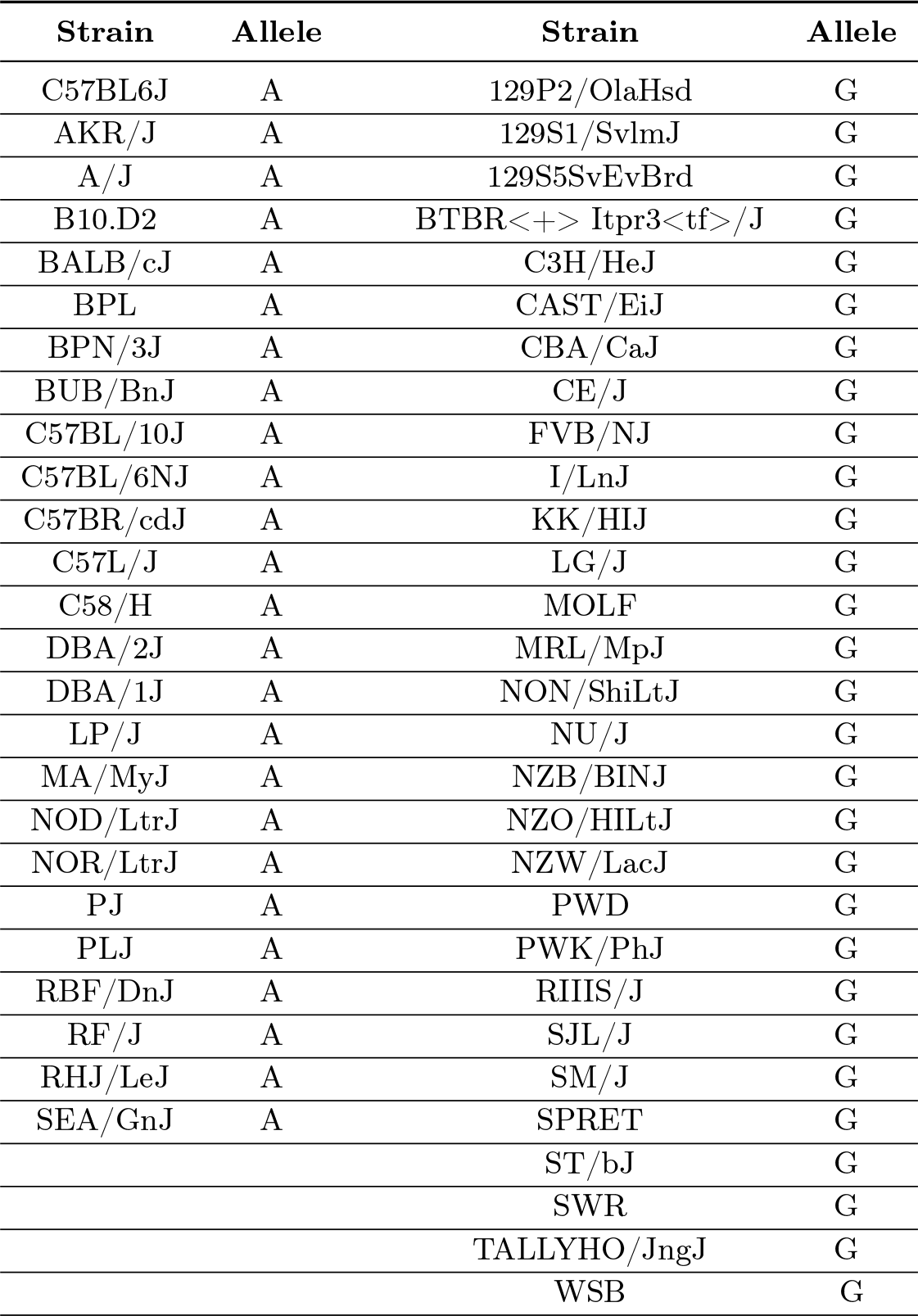
The *Cdh23*^*753(A/G)*^ alleles present in 54 inbred strains are shown. Inbred strains with early onset hearing loss are in bold, and virtually all have the *Cdh23*^*753(A)*^; but many strains with *Cdh23*^*753(A)*^ alleles do not develop early onset hearing loss. I/LnJ mice may have been incorrectly labeled as having an early onset hearing loss in [1]. Examination of I/LnJ data revealed that they were only evaluated at 12 weeks of age, their hearing loss was relatively mild, and there was significant variability in the measured thresholds [1].

| Strain | Allele | Strain | Allele |
| --- | --- | --- | --- |
| C57BL6J | A | 129P2/OlaHsd | G |
| AKR/J | A | 129S1/SvImJ | G |
| A/J | A | 129S5SvEvBrd | G |
| B10.D2 | A | BTBR<+> Itpr3<tf>/J | G |
| BALB/cJ | A | C3H/HeJ | G |
| BPL | A | CAST/EiJ | G |
| BPN/3J | A | CBA/CaJ | G |
| BUB/BnJ | A | CE/J | G |
| C57BL/10J | A | FVB/NJ | G |
| C57BL/6NJ | A | I/LnJ | G |
| C57BR/cdJ | A | KK/HIJ | G |
| C57L/J | A | LG/J | G |
| C58/H | A | MOLF | G |
| DBA/2J | A | MRL/MpJ | G |
| DBA/1J | A | NON/ShiLtJ | G |
| LP/J | A | NU/J | G |
| MA/MyJ | A | NZB/BINJ | G |
| NOD/LtrJ | A | NZO/HILtJ | G |
| NOR/LtrJ | A | NZW/LacJ | G |
| PJ | A | PWD | G |
| PLJ | A | PWK/PhJ | G |
| RBF/DnJ | A | RIIS/J | G |
| RF/J | A | SJL/J | G |
| RHJ/LeJ | A | SM/J | G |
| SEA/GnJ | A | SPRET | G |
|  |  | ST/bJ | G |
|  |  | SWR | G |
|  |  | TALLYHO/JngJ | G |
|  |  | WSB | G |

**Table S2.**
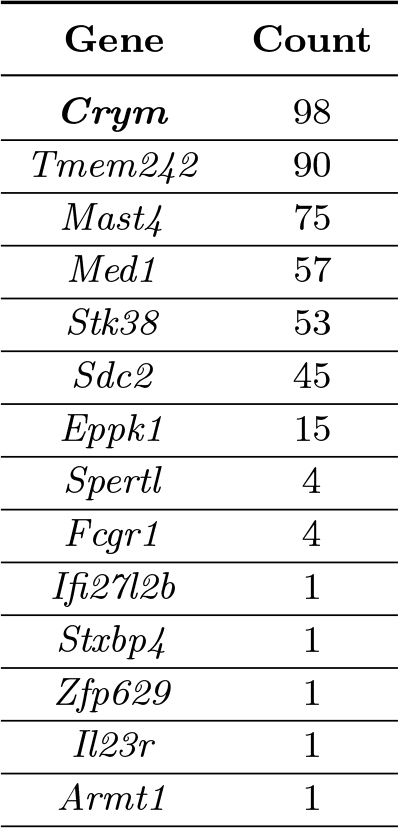
*Crym* was identified by Med-PaLM 2 as the gene whose high impact SNP alleles were most likely to cause spontaneous hearing loss in NOD/LtJ mice. Fourteen genes with high impact SNP alleles were identified by comparing NOD/LtJ genomic sequence with 10 strains that did not manifest age-related hearing loss. Med-PaLM 2 analyzed the gene list 100 times after the gene order was randomly permutated. The number of times each indicated gene was among the top 5 genes identified by Med-PaLM 2 as associated with hearing loss is shown. *Tmem242* was the second most frequently identified gene, but it was not directly associated with hearing loss. *Tmem242* is required for assembly of the mitochondrial ATP synthase complex, and mitochondrial abnormalities (but not *Tmem242* itself) are associated with hearing loss syndrome. The *Med1, Mast4*, and *Stk38* associations resulted from false associations (Mast cells, Med center, etc.) with variants of the gene name.

## Extended data

**Table ED.1.**
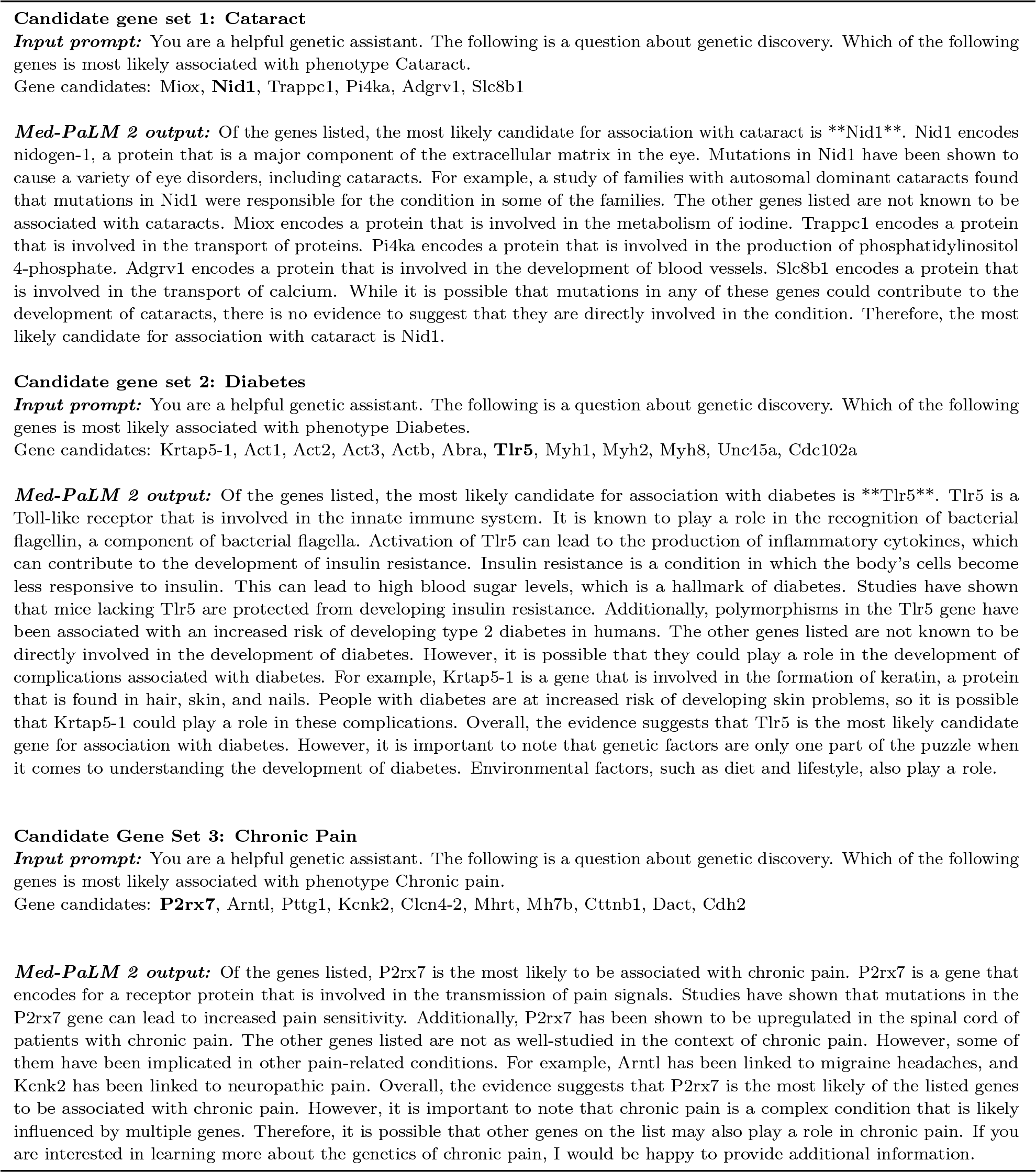
Med-PaLM 2 answers to questions about phenotypes cataract, diabetes, and chronic pain. Candidate gene set 1: Cataract.

**Table ED.2.**
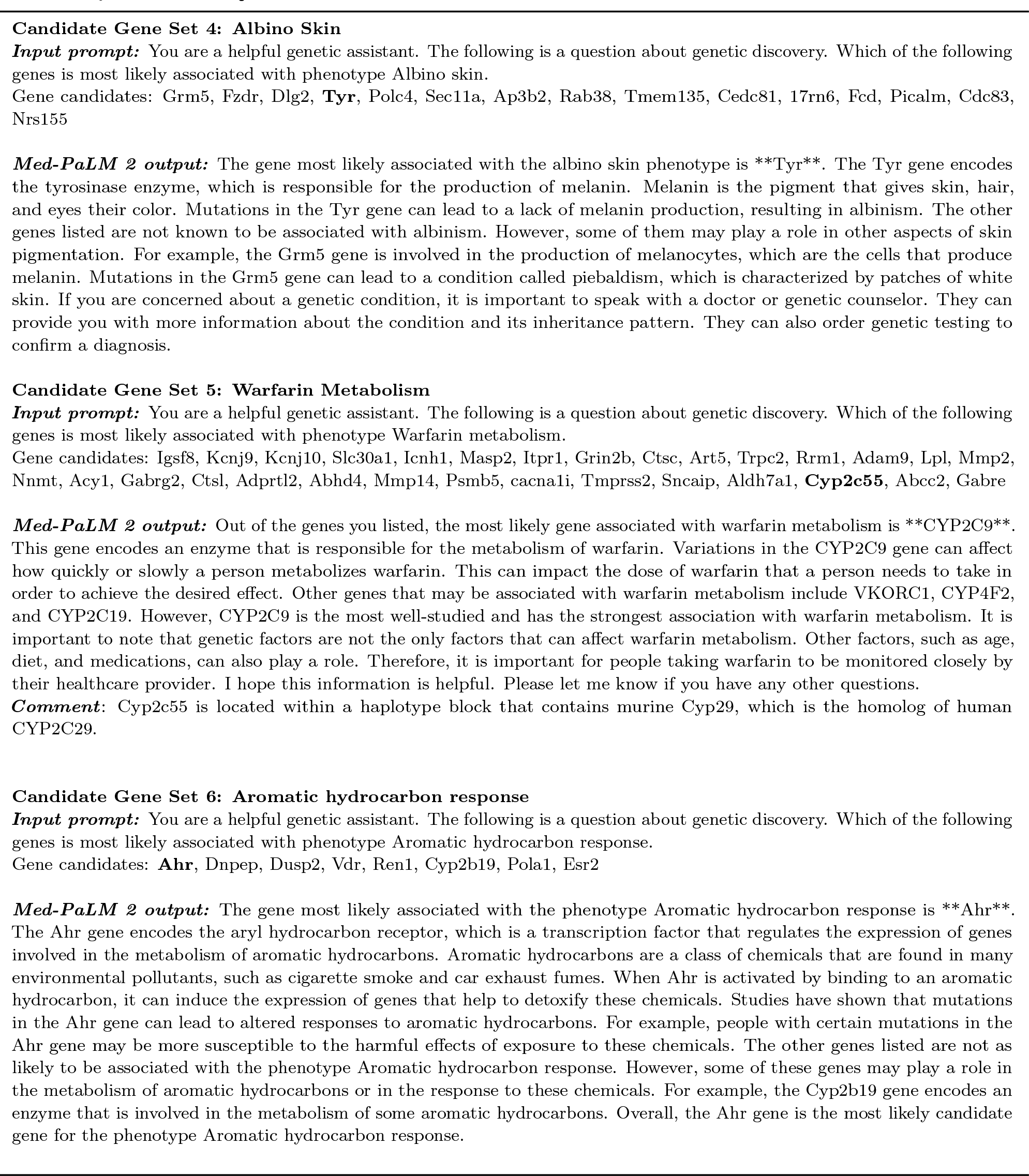
Med-PaLM 2 answers to questions about phenotypes albino skin, warfarin metabolism, and aromatic hydrocarbon response.

**Table ED.3.**
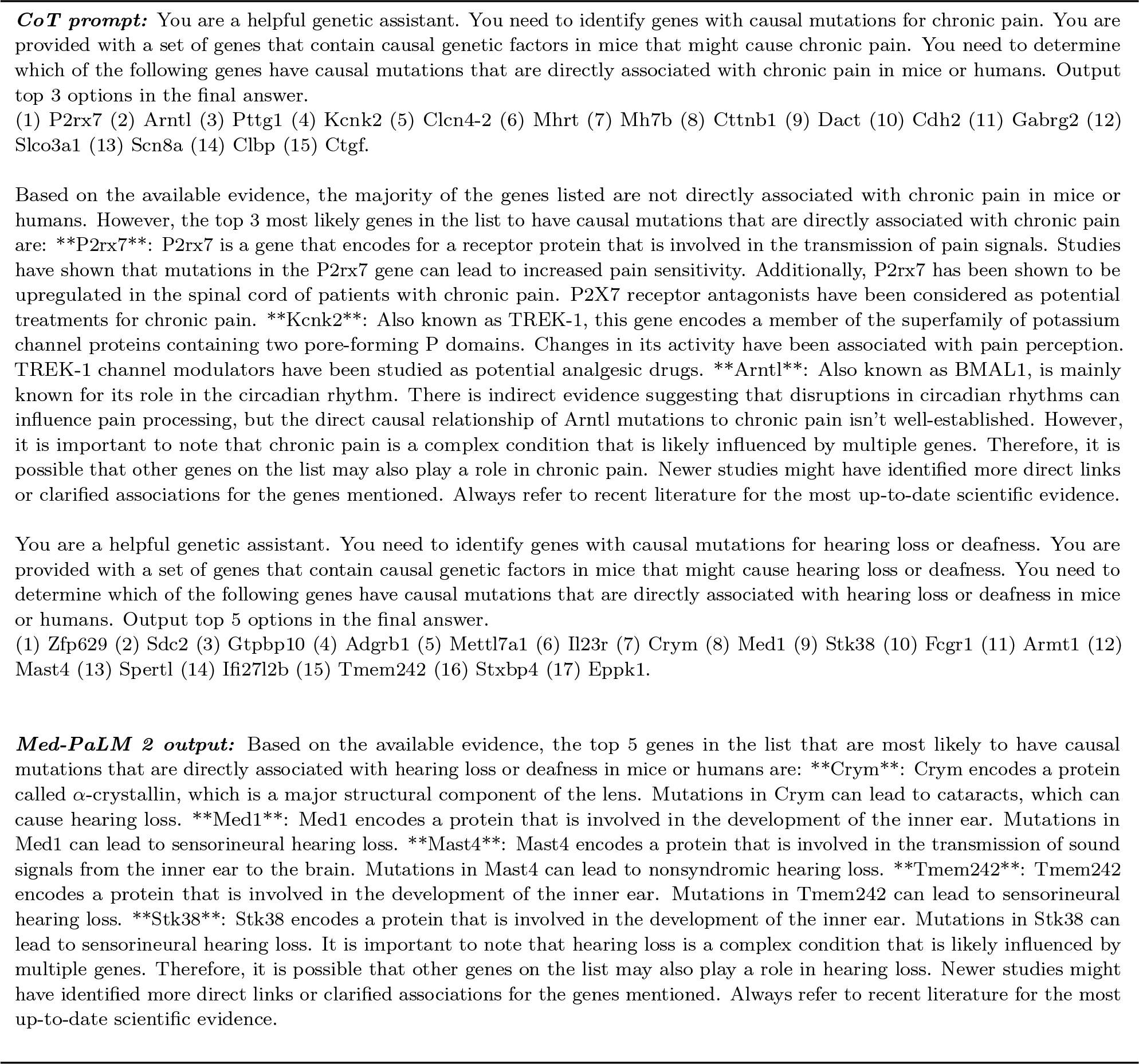
Chain-of-Thought prompt and Med-PaLM 2 answer to the question of identifying 5 genes for hearing loss.

**Table ED.4.**
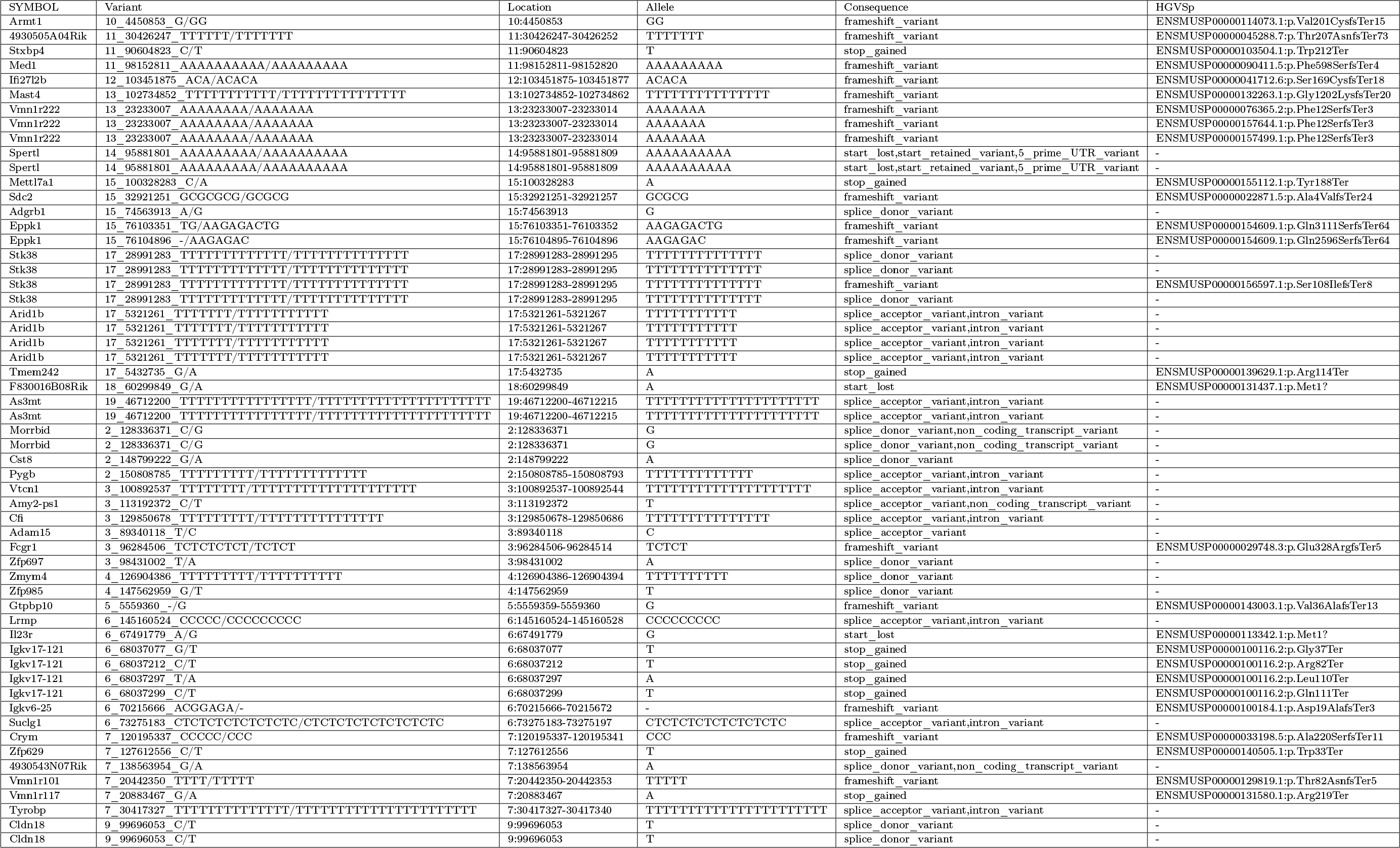
High impact NOD/LtJ-specific SNP alleles.

## Notes

### Competing Interest Statement

The Stanford University Medical School authors have no competing interests. Tao Tu, Vivek Natarajan, and Anil Palepu are employees of Alphabet and may own stock as part of their standard compensation package.

## References

1. Fang, Z. & Peltz, G. An automated multi-modal graph-based pipeline for mouse genetic discovery. Bioinformatics 38, 3385–3394 (2022).

2. Vaswani, A., Shazeer, N., Parmar, N., Uszkoreit, J., Jones, L., Gomez, A. N., Kaiser, Ł. & Polosukhin, I. Attention is all you need. Advances in neural information processing systems 30 (2017).

3. Chowdhery, A., Narang, S., Devlin, J., Bosma, M., Mishra, G., Roberts, A., Barham, P., Chung, H. W., Sutton, C., Gehrmann, S., et al. Palm: Scaling language modeling with pathways. arXiv preprint arXiv:2204.02311 (2022).

4. Singhal, K., Azizi, S., Tu, T., Mahdavi, S. S., Wei, J., Chung, H. W., Scales, N., Tanwani, A., Cole-Lewis, H., Pfohl, S., et al. Large language models encode clinical knowledge. Nature 620, 172–180 (2023).

5. Singhal, K., Tu, T., Gottweis, J., Sayres, R., Wulczyn, E., Hou, L., Clark, K., Pfohl, S., Cole-Lewis, H., Neal, D., et al. Towards expert-level medical question answering with large language models. arXiv preprint arXiv:2305.09617 (2023).

6. Liao, G., Wang, J., Guo, J., Allard, J., Cheng, J., Ng, A., Shafer, S., Puech, A., McPherson, J. D., Foernzler, D., et al. In silico genetics: identification of a functional element regulating H2-E α gene expression. Science 306, 690–695 (2004).

7. Guo, Y., Weller, P., Farrell, E., Cheung, P., Fitch, B., Clark, D., Wu, S.-y., Wang, J., Liao, G., Zhang, Z., et al. In silico pharmacogenetics of warfarin metabolism. Nature biotechnology 24, 531–536 (2006).

8. Sorge, R. E., Trang, T., Dorfman, R., Smith, S. B., Beggs, S., Ritchie, J., Austin, J.-S., Zaykin, D. V., Meulen, H. V., Costigan, M., et al. Genetically determined P2X7 receptor pore formation regulates variability in chronic pain sensitivity. Nature medicine 18, 595–599 (2012).

9. Smith, P. A. K+ channels in primary afferents and their role in nerve injury-induced pain. Frontiers in Cellular Neuroscience 14, 566418 (2020).

10. Cregg, R., Momin, A., Rugiero, F., Wood, J. N. & Zhao, J. Pain channelopathies. The Journal of physiology 588, 1897–1904 (2010).

11. Alloui, A., Zimmermann, K., Mamet, J., Duprat, F., Noël, J., Chemin, J., Guy, N., Blondeau, N., Voilley, N., Rubat-Coudert, C., et al. TREK-1, a K+ channel involved in polymodal pain perception. The EMBO journal 25, 2368–2376 (2006).

12. Takahashi, J. S. Transcriptional architecture of the mammalian circadian clock. Nature Reviews Genetics 18, 164–179 (2017).

13. Bumgarner, J. R., Walker II, W. H. & Nelson, R. J. Circadian rhythms and pain. Neuroscience & Biobehavioral Reviews 129, 296–306 (2021).

14. Zheng, Q. Y., Johnson, K. R. & Erway, L. C. Assessment of hearing in 80 inbred strains of mice by ABR threshold analyses. Hearing research 130, 94–107 (1999).

15. Noben-Trauth, K., Zheng, Q. Y. & Johnson, K. R. Association of cadherin 23 with polygenic inheritance and genetic modification of sensorineural hearing loss. Nature genetics 35, 21–23 (2003).

16. Boëda, B., El-Amraoui, A., Bahloul, A., Goodyear, R., Daviet, L., Blanchard, S., Perfettini, I., Fath, K. R., Shorte, S., Reiners, J., et al. Myosin VIIa, harmonin and cadherin 23, three Usher I gene products that cooperate to shape the sensory hair cell bundle. The EMBO journal 21, 6689–6699 (2002).

17. Siemens, J., Kazmierczak, P., Reynolds, A., Sticker, M., Littlewood-Evans, A. & Müller, U. The Usher syndrome proteins cadherin 23 and harmonin form a complex by means of PDZ-domain interactions. Proceedings of the National Academy of Sciences 99, 14946–14951 (2002).

18. Arslan, A., Guan, Y., Fang, Z., Chen, X., Donaldson, R., Zhu, W., Ford, M., Wu, M., Zheng, M., Dill, D. L., et al. High throughput computational mouse genetic analysis. BioRxiv, 2020–09 (2020).

19. Arslan, A., Fang, Z., Wang, M., Cheng, Z., Yoo, B., Bejerano, G. & Peltz, G. Analysis of structural variation among inbred mouse strains identifies genetic factors for autism-related traits. BioRxiv, 2021–02 (2021).

20. Arslan, A., Fang, Z., Wang, M., Tan, Y., Cheng, Z., Chen, X., Guan, Y., J. Pisani, L., Yoo, B., Bejerano G., et al. Analysis of structural variation among inbred mouse strains. BMC genomics 24, 97 (2023).

21. Prochazka, M., Serreze, D. V., Frankel, W. N. & Leiter, E. H. NOR/Lt mice: MHC-matched diabetes-resistant control strain for NOD mice. Diabetes 41, 98–106 (1992).

22. Yoshimura, H., Takumi, Y., Nishio, S.-y., Suzuki, N., Iwasa, Y.-i. & Usami, S.-i. Deafness gene expression patterns in the mouse cochlea found by microarray analysis. PLoS One 9, e92547 (2014).

23. Abe, S., Katagiri, T., Saito-Hisaminato, A., Usami, S.-i., Inoue, Y., Tsunoda, T. & Nakamura, Y. Identification of CRYM as a candidate responsible for nonsyndromic deafness, through cDNA microarray analysis of human cochlear and vestibular tissues. The American Journal of Human Genetics 72, 73–82 (2003).

24. Oshima, A., Suzuki, S., Takumi, Y., Hashizume, K., Abe, S. & Usami, S. CRYM mutations cause deafness through thyroid hormone binding properties in the fibrocytes of the cochlea. Journal of medical genetics 43, e25–e25 (2006).

25. Kinney, C. J. & Bloch, R. J. µ-Crystallin: A thyroid hormone binding protein. Endocrine regulations 55, 89–102 (2021).

26. Aksoy, O., Hantusch, B. & Kenner, L. Emerging role of T3-binding protein µ-crystallin (CRYM) in health and disease. Trends in Endocrinology & Metabolism (2022).

27. Borel, F., Hachi, I., Palencia, A., Gaillard, M.-C. & Ferrer, J.-L. Crystal structure of mouse mu-crystallin complexed with NADPH and the T3 thyroid hormone. The FEBS journal 281, 1598–1612 (2014).

28. Hou, X., Yasuda, S. P., Yamaguchi, M., Suzuki, S., Seki, Y., Ouchi, T., Mao, T., Prakhongcheep, O., Shitara, H. & Kikkawa, Y. Impacts of an age-related hearing loss allele of cadherin 23 on severity of hearing loss in ICR and NOD/Shi mice. Biochemical and Biophysical Research Communications 674, 147–153 (2023).

29. Suzuki, S., Suzuki, N., Mori, J.-i., Oshima, A., Usami, S. & Hashizume, K. µ-Crystallin as an intracellular 3, 5, 3-triiodothyronine holder in vivo. Molecular endocrinology 21, 885–894 (2007).

30. Church, G. Compelling reasons for repairing human germlines. N Engl J Med 377, 1909–1911 (2017).

31. Yang, Y., Muzny, D. M., Reid, J. G., Bainbridge, M. N., Willis, A., Ward, P. A., Braxton, A., Beuten, J., Xia, F., Niu, Z., et al. Clinical whole-exome sequencing for the diagnosis of mendelian disorders. New England Journal of Medicine 369, 1502–1511 (2013).

32. Lee, H., Deignan, J. L., Dorrani, N., Strom, S. P., Kantarci, S., Quintero-Rivera, F., Das, K., Toy, T., Harry, B., Yourshaw, M., et al. Clinical exome sequencing for genetic identification of rare Mendelian disorders. Jama 312, 1880–1887 (2014).

33. Ng, S. B., Buckingham, K. J., Lee, C., Bigham, A. W., Tabor, H. K., Dent, K. M., Huff, C. D., Shannon, P. T., Jabs, E. W., Nickerson, D. A., et al. Exome sequencing identifies the cause of a mendelian disorder. Nature genetics 42, 30–35 (2010).

34. Ng, S. B., Turner, E. H., Robertson, P. D., Flygare, S. D., Bigham, A. W., Lee, C., Shaffer, T., Wong, M., Bhattacharjee, A., Eichler, E. E., et al. Targeted capture and massively parallel sequencing of 12 human exomes. Nature 461, 272–276 (2009).

35. Dragojlovic, N., Elliott, A. M., Adam, S., van Karnebeek, C., Lehman, A., Mwenifumbo, J. C., Nelson, T. N., du Souich, C., Friedman, J. M. & Lynd, L. D. The cost and diagnostic yield of exome sequencing for children with suspected genetic disorders: a benchmarking study. Genetics in Medicine 20, 1–9 (2018).

36. Wang, K., Li, M. & Hakonarson, H. ANNOVAR: functional annotation of genetic variants from high-throughput sequencing data. Nucleic acids research 38, e164–e164 (2010).

37. Jagadeesh, K. A., Wenger, A. M., Berger, M. J., Guturu, H., Stenson, P. D., Cooper, D. N., Bernstein, J. A. & Bejerano, G. M-CAP eliminates a majority of variants of uncertain significance in clinical exomes at high sensitivity. Nature genetics 48, 1581–1586 (2016).

38. Jagadeesh, K. A., Paggi, J. M., Ye, J. S., Stenson, P. D., Cooper, D. N., Bernstein, J. A. & Bejerano, G. S-CAP extends pathogenicity prediction to genetic variants that affect RNA splicing. Nature genetics 51, 755–763 (2019).

39. Singleton, M. V., Guthery, S. L., Voelkerding, K. V., Chen, K., Kennedy, B., Margraf, R. L., Durtschi, J., Eilbeck, K., Reese, M. G., Jorde, L. B., et al. Phevor combines multiple biomedical ontologies for accurate identification of disease-causing alleles in single individuals and small nuclear families. The American Journal of Human Genetics 94, 599–610 (2014).

40. Smedley, D., Jacobsen, J. O., Jäger, M., Köhler, S., Holtgrewe, M., Schubach, M., Siragusa, E., Zemojtel, T., Buske, O. J., Washington, N. L., et al. Next-generation diagnostics and disease-gene discovery with the Exomiser. Nature protocols 10, 2004–2015 (2015).

41. Jagadeesh, K. A., Birgmeier, J., Guturu, H., Deisseroth, C. A., Wenger, A. M., Bernstein, J. A. & Bejerano, G. Phrank measures phenotype sets similarity to greatly improve Mendelian diagnostic disease prioritization. Genetics in Medicine 21, 464–470 (2019).

42. Birgmeier, J., Haeussler, M., Deisseroth, C. A., Steinberg, E. H., Jagadeesh, K. A., Ratner, A. J., Guturu, H., Wenger, A. M., Diekhans, M. E., Stenson, P. D., et al. AMELIE speeds Mendelian diagnosis by matching patient phenotype and genotype to primary literature. Science translational medicine 12, eaau9113 (2020).

43. Caro-Llopis, A., Rosello, M., Orellana, C., Oltra, S., Monfort, S., Mayo, S. & Martinez, F. De novo mutations in genes of mediator complex causing syndromic intellectual disability: mediatorpathy or transcriptomopathy? Pediatric research 80, 809–815 (2016).

44. Rohowetz, L. J., Mardelli, M. E., Duncan, R. S., Riordan, S. M. & Koulen, P. The contribution of anterior segment abnormalities to changes in intraocular pressure in the DBA/2J mouse model of glaucoma: DBA/2J-Gpnmb+/SjJ mice as critical controls. Frontiers in Neuroscience 15, 801184 (2022).

45. Cheng, J., Novati, G., Pan, J., Bycroft, C., Žemgulytė, A., Applebaum, T., Pritzel, A., Wong, L. H., Zielinski, M., Sargeant, T., et al. Accurate proteome-wide missense variant effect prediction with AlphaMissense. Science 381, eadg7492 (2023).

## References

46. Kilkenny, C., Browne, W. J., Cuthill, I. C., Emerson, M. & Altman, D. G. Improving bioscience research reporting: the ARRIVE guidelines for reporting animal research. Journal of Pharmacology and Pharmacotherapeutics 1, 94–99 (2010).

47. Seist, R., Landegger, L. D., Robertson, N. G., Vasilijic, S., Morton, C. C. & Stankovic, K. M. Cochlin deficiency protects against noise-induced hearing loss. Frontiers in Molecular Neuroscience 14, 670013 (2021).

48. Early, S., Saad, M. A., Mallidi, S., Mansour, A., Seist, R., Hasan, T. & Stankovic, K. M. A fluorescent photoimmunoconjugate for imaging of cholesteatoma. Scientific Reports 12, 19905 (2022).

49. Chen, J., Landegger, L. D., Sun, Y., Ren, J., Maimon, N., Wu, L., Ng, M. R., Chen, J. W., Zhang, N., Zhao, Y., et al. A cerebellopontine angle mouse model for the investigation of tumor biology, hearing, and neurological function in NF2-related vestibular schwannoma. Nature protocols 14, 541–555 (2019).

50. Li, H. & Durbin, R. Fast and accurate short read alignment with Burrows–Wheeler transform. bioinformatics 25, 1754–1760 (2009).

51. Li, H., Handsaker, B., Wysoker, A., Fennell, T., Ruan, J., Homer, N., Marth, G., Abecasis, G., Durbin, R. & Subgroup, G. P. D. P. The sequence alignment/map format and SAMtools. bioinformatics 25, 2078–2079 (2009).

52. McLaren, W., Gil, L., Hunt, S. E., Riat, H. S., Ritchie, G. R., Thormann, A., Flicek, P. & Cunningham, F. The ensembl variant effect predictor. Genome biology 17, 1–14 (2016).

